# PHYTOCHROME C regulation of *PHOTOPERIOD1* is mediated by *EARLY FLOWERING 3* in *Brachypodium distachyon*

**DOI:** 10.1101/2022.10.11.511813

**Authors:** Daniel P. Woods, Weiya Li, Richard Sibout, Mingqin Shao, Debbie Laudencia-Chingcuanco, John P. Vogel, Jorge Dubcovsky, Richard M. Amasino

## Abstract

Daylength sensing in many plants is critical for coinciding the timing of flowering with the appropriate season. Temperate-climate-adapted grasses such as *Brachypodium distachyon* flower during the spring when days are becoming longer. The photoreceptor PHYTOCHROME C is essential for long-day (LD) flowering in *B. distachyon. PHYC* is required for the LD activation of a suite of genes in the photoperiod pathway including *PHOTOPERIOD1* (*PPD1*) that, in turn, result in the activation of *FLOWERING LOCUS T* (*FT1*)/*FLORIGEN*, which causes flowering. Thus, *phyC* mutants are extremely delayed in flowering. Here we show that PHYC-mediated activation of PPD1 occurs via *EARLY FLOWERING 3* (*ELF3*), a component of the evening complex in the circadian clock. The extreme delay of flowering of the *phyC* mutant disappears when combined with an *elf3* loss-of-function mutation. Moreover, the dampened *PPD1* expression in *phyC* mutant plants is elevated in *phyC/elf3* mutant plants consistent with the rapid flowering of the double mutant. We show that loss of *PPD1* function also results in reduced *FT1* expression levels and extremely delayed flowering consistent with reports from wheat and barley. Additionally, *elf3* mutant plants have elevated expression levels of *PPD1* and we show that overexpression of *ELF3* results in delayed flowering, which is associated with a reduction of *PPD1* and *FT1*, demonstrating ELF3 represses *PPD1* transcription, consistent with previous studies showing that ELF3 binds to the *PPD1* promoter. Indeed, *PPD1* is the main target of ELF3-mediated flowering as *elf3/ppd1* double mutant plants are delayed flowering. Our results indicate that *ELF3* operates downstream from *PHYC* and acts as a repressor of *PPD1* in the photoperiod flowering pathway of *B. distachyon*.

**AUTHOR SUMMARY:** Daylength is an important environmental cue that plants and animals use to coincide important life history events with a proper season. In plants, timing of flowering to a particular season is an essential adaptation to many ecological niches. Perceiving changes in daylength starts with the perception of light via specific photoreceptors such as phytochromes. In temperate grasses, how daylength perception is integrated into downstream pathways to trigger flowering is not fully understood. However, some of the components involved in the translation of daylength perception into the induction of flowering in temperate grasses have been identified from studies of natural variation. For example, specific alleles of two genes called *EARLY FLOWERING 3* (*ELF3*) and *PHOTOPERIOD1* (*PPD1*) have been selected during breeding of different wheat and barley varieties to modulate the photoperiodic response to maximize reproduction in different environments. Here, we show in the temperate grass model *Brachypodium distachyon* that the translation of the light signal perceived by phytochromes into a flowering response is mediated by *ELF3*, and that *PPD1* is genetically downstream of *ELF3* in the photoperiodic flowering pathway. These results provide a genetic framework for understanding the photoperiodic response in temperate grasses that include agronomically important crops such as wheat, oats, barley, and rye.

## INTRODUCTION

The transition from vegetative to reproductive development is an important developmental decision for which the timing is often directly influenced by the environment (e.g. [1–4]). This critical life history trait has been shaped over evolutionary time to enable reproduction to coincide with the particular time of year that is most favorable for flower and seed development. Moreover, breeding to adjust the timing of the flowering transition in crops has been critical for adapting various crop varieties to changing environments and to increase yield e.g. [5].

In many plant species, changes in day-length and/or temperature provide the seasonal cues that result in flowering occurring during a specific time of year [1,6], a response known as photoperiodism [1]. Many temperate grasses such as *Brachypodium distachyon* (*B. distachyon*), wheat, and barley that flower in the spring or early summer months in response to increasing day-lengths, and thus are referred to as long-day (LD) plants [7]. *B. distachyon* is closely related to the core pooid clade comprising wheat, oats, barley, and rye and has a number of attributes that make it an attractive grass model organism suitable for developmental genetics research [8,9]. Unlike temperate grasses, many tropical grasses such as rice, flower when the days become shorter and thus are referred to as short-day (SD) plants [6].

Variation in the LD promotion of flowering in temperate grasses such as wheat and barley are often due to allelic variation at *PHOTOPERIOD1* (*PPD1*), a member of the pseudo-response regulator (PRR) gene family, hence *PPD1* is also known as *PSEUDO RESPONSE REGULATOR 37* (*PRR37*) [10, 11]. Natural variation in *PPD1* resulting in either hypomorphic alleles as found in barley or dominant *PPD1* alleles as found in tetraploid or hexaploid wheat impact flowering [10,11–15]. Specifically, natural recessive mutations in the conserved CONSTANS, CONSTANS-LIKE and TIMING OF CAB EXPRESSION 1 (CCT) putative DNA binding domain in the barley PPD1 protein cause photoperiod insensitivity and delayed flowering under LD [11,12], whereas wheat photoperiod insensitivity is linked to overlapping large deletions in the promoter region of *PPD1* in either the A [13] or D genome homeologs [10]. These large deletions result in elevated expression of *PPD1*, particularly during dawn, causing rapid flowering even under non-inductive SD conditions [13]. It is worth noting that although these wheat lines are referred to as photoperiod insensitive (PI) varieties they still flower earlier under LD than under SD if the timing of flowering is measured as the emergence of the wheat spike (heading time) [16]. It has been hypothesized that the large deletion within the *PPD1* promoter might remove a binding site for one or more transcriptional repressors [13]. To date, natural variation studies of flowering in *B. distachyon* have not pointed to allelic variation at *PPD1* and thus its role in LD flowering in *B. distachyon* is not known [17–21].

Variation in *EARLY FLOWERING 3* (*ELF3*) also impacts photoperiodic flowering in grasses, including wheat [22,23], barley [24,25], and rice [26]. In these plants, natural variation in *ELF3* allows growth at latitudes that otherwise would not be inductive for flowering, enabling these crops to be grown in regions with short growing seasons [5]. For example, *early maturity* (*eam*) loci have been used by breeders to allow barley to grow at higher latitudes in regions of northern Europe with short growing seasons [24,27]. The *eam8* mutant in the barley ortholog of *ELF3*, is a loss-of-function mutation that accelerates flowering under SD or LDs [24,25] similar to *elf3* loss-of-function alleles described previously in the eudicot model *Arabidopsis thaliana* (*A. thaliana*) [28]. Moreover, loss of function of *ELF3* in *B. distachyon* also results in rapid flowering under SD and LD, and expression of the *B. distachyon* ELF3 protein is able to rescue the *A. thaliana elf3* mutant, demonstrating a conserved role of *ELF3* in flowering across angiosperm diversification [29–31].

Work in *A. thaliana* has shown that *ELF3* is an important component of the circadian clock that acts as a bridge protein within a trimeric protein complex that also contains *LUX ARRTHYHMO* (LUX), and *EARLY FLOWERING 4* (*ELF4*) and is referred to as the evening complex (EC) [32]. Loss-of-function mutations in any of the proteins that make up the EC results in rapid flowering and disrupted clock function [33–36]. The peak expression of the EC at dusk is involved in the direct transcriptional repression of genes that make up the morning loop of the circadian clock including *A. thaliana PRR7* and *PRR9*, which are paralogs of grass *PPD1* and *PRR73* [37–39]. Recently, it has been shown that the EC also directly represses *PRR37, PRR95*, and *PRR73* in rice; *PRR37* is the rice ortholog of *PPD1*, again suggesting at least in part conservation of the role of the EC across flowering plant diversification [40]. Furthermore, *elf3* mutants in barley, wheat, and *B. distachyon* have elevated *PPD1* expression [23,24,30] indicating *ELF3* may impact flowering in part via *PPD1*, but to what extent remains to be determined.

The photoperiod and circadian pathways converge in the transcriptional activation of florigen/*FLOWERING LOCUS1* (*FT1*) in leaves [6,41]. In temperate grasses, *PPD1* is required for the LD induction of *FT1*, whereas in *A. thaliana CONSTANS* (*CO*) is the main photoperiodic gene required for *FT1* activation in LD [16,42,43]. There are two *CO*-like genes in temperate grasses, interestingly, in the presence of functional *PPD1, co1co2* wheat plants have a modest earlier heading phenotype suggesting they are in fact mild floral repressors, but in the absence of *PPD1 CO1* acts as a flowering promoter under LD [16]. To date, no null *co1co2* double mutants have been reported in *B. distachyon*. However, RNAi knock-down of *co1* does result in a 30-day delay in flowering under 16h LD [44] and overexpression of *CO1* leads to earlier flowering in SD [44]. These results indicate that in *B. distachyon CO1* does have a promoting role in flowering even in the presence of a functional *PPD1* gene, and suggest potential differences in the role of *CO1* in the regulation of flowering between *B. distachyon* and wheat.

Once FT1 is activated by LD it interacts with the bZIP transcription factor FD which triggers the expression of the MADS-box transcription factor *VERNALIZATION1* (*VRN1*) [45,46]. VRN1 in turn upregulates the expression of *FT1* forming a positive feedback loop that overcomes the repression from the zinc finger and CCT domain-containing transcription factor *VERNALIZATION2* (*VRN2*) [17,47–50, 84]. The FT1 protein is then thought to migrate from the leaves to the shoot apical meristem, as shown in *A. thaliana* and rice [51,52], to induce the expression of floral homeotic genes including *VRN1*, thus converting the vegetative meristem to a floral meristem under favorable LD photoperiods.

Light signals are perceived initially by photoreceptors that initiate a signal transduction cascade impacting a variety of different outputs including developmental responses to light [53]. The sensing of light is accomplished by complementary photoreceptors: phytochromes which measure the ratio of red and far-red light whereas cryptochromes and phototropins detect blue light frequencies [54,55]. The phytochromes form homodimers that upon exposure of plants to red light undergo a confirmation shift to an active form causing the activation of a suite of downstream genes [54]. Exposure of plants to far-red or dark conditions causes photo-reversion of the phytochromes to an inactive state [54].

There are three phytochromes in temperate grasses referred to as PHYTOCHROME A (PHYA), PHYTOCHROME B (PHYB), and PHYTOCHROME C (PHYC) [56]. Functional analyses of these phytochromes in temperate grasses revealed that PHYB and PHYC play a major role in the LD induction of flowering because loss-of-function mutations in both of these genes results in extremely delayed flowering [57–59] whereas loss-of-function mutations in *PHYA* in *B. distachyon* results in only a modest delay of flowering under inductive LD, indicating PHYB and PHYC are the main light receptors required for photoperiodic flowering in temperate grasses [29].The important role of PHYC in photoperiodic flowering is not universal as loss of *phyC* function in *A. thaliana* and rice only has small effects on flowering [60,61].

In temperate grasses, PHYB and PHYC are required for the transcriptional activation of a suite of genes involved in the photoperiod pathway, including *PPD1, CO1*, and FT1, and ectopic expression of *FT1* in the *B. distachyon phyC* background results in rapid flowering—a reversal of the *phyC* single-mutant phenotype [57,58,62]. Moreover, consistent with PHYB/C acting at the beginning of the photoperiodic flowering signal cascade, expression of genes encoding components of the circadian clock are also severely dampened in the *phyB* and *phyC* mutant backgrounds [29,57–59]. An exception to this is that the expression of *ELF3* is not altered in the temperate grass phytochrome mutants [29,58–59]. Recently, in *B. distachyon* it has been shown that PHYC can interact with ELF3, and this interaction destabilizes the ELF3 protein indicating that the regulation of ELF3 by PHY is at least in part at the protein level consistent with previous studies from *A. thaliana*, rice and the companion study in wheat [29,40,63, 68]. At present it is not clear to what extent the regulation of ELF3 by PHYs is critical for photoperiodic flowering.

Here, we show in *B. distachyon* by analyzing *phyC/efl3* double mutant plants that indeed the light signal perceived by phytochromes is mediated through ELF3 for photoperiodic flowering. The extreme delayed flowering of the *phyC* mutant disappears in the *phyC/elf3* double mutant plants which flower as rapidly as *elf3* single mutant plants. Moreover, the expression profiles of genes in the photoperiod pathway are similar between *elf3* and *phyC/elf3* mutant plants compared to *phyC* mutant plants. Thus, *elf3* is completely epistatic to *phyC* at the phenotypic and molecular levels. Furthermore, we show strong, environment-dependent genetic interactions between *ELF3* and *PPD1*, which indicates that *PPD1* is a main target of ELF3-mediated repression of flowering. These results provide a genetic and molecular framework to understand photoperiodic flowering in the temperate grasses.

## RESULTS

### The rapid flowering *elf3* mutant is epistatic to the delayed flowering *phyC* mutant

Previous studies in *B. distachyon* showed that PHYC can affect the stability of the ELF3 protein, and that the transcriptome of a *phyC* mutant resembles that of a plant with elevated ELF3 signaling [29]. Thus, it has been suggested that the extreme delayed flowering phenotype of the *phyC* mutant [58] could be mediated by ELF3 [29]. To test the extent to which the translation of the light signal perceived by PHYC to control flowering is mediated by ELF3, we generated *elf3/phyC* double mutant plants and evaluated the flowering of the double mutant relative to that of *elf3* and *phyC* single mutants as well as Bd21-3 wild type plants under 16h-LD and 8h-SD (Fig 1).

**Fig 1.**
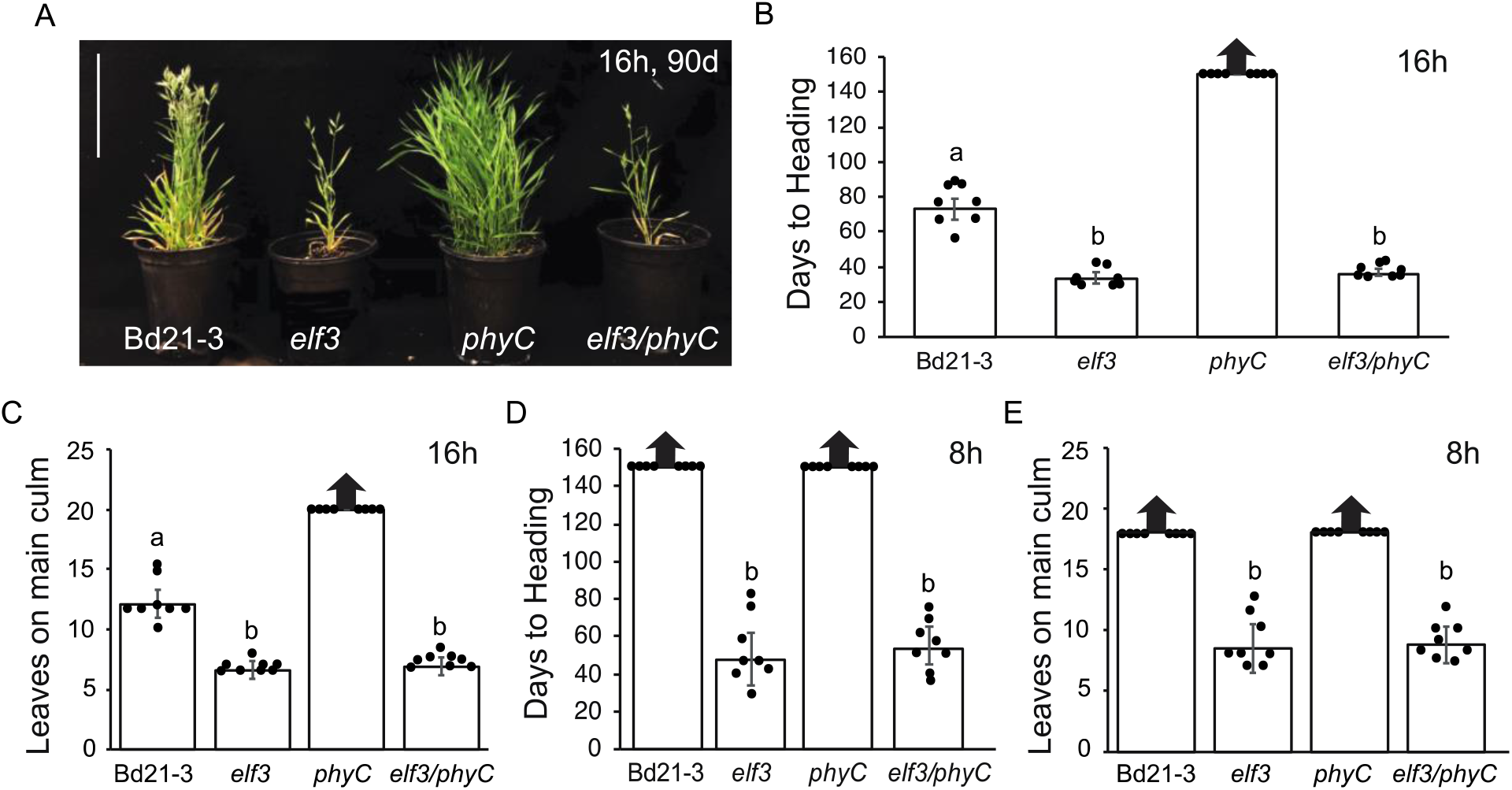
The rapid flowering of the *elf3* mutant is epistatic to the delayed flowering of the *phyC* mutant (**A**) Representative images of Bd21-3 wild-type, *elf3* mutant, *phyC* mutant and *elf3/phyC* double mutant plants grown in a 16h photoperiod at 90d after germination. Bar=17cm. (**B, D**) Flowering times under 16h (**B**) or 8h daylengths (**D**) measured as days to heading of Bd21-3, *elf3, phyC*, and *elf3/phyC*. (**C**) Flowering phenotypes under 16h (**C**) or 8h daylengths (**E**) measured as the number of leaves on the parent culm at time of heading. Bars represent the average of 8 plants +/- SD. Arrows above bars indicate that none of the plants flowered at the end of the experiment (150d). Letters (a, b) indicate statistical differences (*p* < 0.05) according to a Tukey’s HSD test used to perform multiple comparisons.

Under either 16h LD or 8 SD photoperiods, we found that *elf3* is epistatic to *phyC*. Specifically, in LD *elf3/phyC* double mutant plants flowered rapidly by 38 days with 6.9 leaves similar to *elf3* mutant plants that flowered by 34 days with 6.6 leaves (Fig 1A, B). In contrast, *phyC* mutant plants had not flowered after 150 days with greater than 20 leaves when the experiment was terminated, and Bd21-3 wild-type plants flower by 72 days with 12 leaves consistent with previous studies [49,58]. In 8h SD, *elf3/phyC* double mutant plants also flowered rapidly by 54 days with 8.8 leaves similar to *elf3* mutant plants that flowered by 48 days with 8.5 leaves (Fig 1D, E). In contrast, both Bd21-3 wild-type and *phyC* mutants had not flowered by 150 days with >18 leaves when the experiment was terminated (Fig 1D, E). These results indicate that the extreme delayed flowering mutant phenotype of *phyC* in *B. distachyon* is mediated by ELF3.

### Effect of mutations in *PHYC* and *ELF3* on the transcriptional profiles of flowering time genes

To further understand how *PHYC* and *ELF3* affect flowering at a molecular level, we compared the mRNA levels of *B. distachyon* orthologs of the photoperiodic and vernalization genes *FT1, VRN1, PPD1, VRN2, CO1*, and *CO2* across a diurnal cycle in 16h LD in the *phyC* and *elf3* single mutants versus the *elf3/phyC* double mutant (Fig. 2). We were particularly interested in determining how the expression profiles of “flowering-time genes” in the *elf3/phyC* double mutant compared to the *elf3* and *phyC* single mutant expression profiles. Consistent with the rapid flowering of the *elf3* and *elf3/phyC* mutants, the mRNA expression levels of *FT1* and *VRN1* in these lines are significantly higher than the levels in wild-type and phyC mutant plants across all the time points tested (Fig 2 A, D). Moreover, the overall expression profiles of *FT1* and *VRN1* between the *elf3* and *elf3/phyC* mutant plants were similar throughout the day. This is in contrast with the *phyC* mutant, in which *FT1* and *VRN1* mRNA levels were lower than wild-type throughout the day consistent with the delayed flowering phenotype of *phyC*. Despite the elevated levels of *FT1* and *VRN1* in both *elf3* and *elf3/phyC* consistent with their rapid flowering, the expression of the floral repressor, *VRN2*, has a similar elevated expression profile throughout the day in both *elf3* and *elf3/phyC* relative to wild-type or *phyC* single-mutant plants (Fig 2E). The elevated *VRN2* expression levels in *elf3* mutant plants are consistent with previous results in *B. distachyon* and other grasses [29, 30, 40, 64, 65]. The transcriptional profile of *CO1* was similar in both the *elf3* and *elf3/phyC* mutant plants with elevated expression compared to wild-type between zt4-8 and then lower than wild-type between zt12-20 (Fig 2C). A similar expression pattern was found for *Hd1* (rice *CO* homolog) in the *elf3-1/elf3-2* double mutant in rice [40]. Consistent with previous reports, *CO1* expression levels remained low in *Brachypodium phyC* mutants throughout a diurnal cycle [58]. By contrast *CO1* expression is significantly upregulated in the *phyC* mutants in wheat [57] indicating another difference in the regulation of *CO1* between these two species. Lastly, the *CO2* expression profiles were similar between wild-type, *elf3*, and *elf3/phyC* whereas *CO2* mRNA levels were lower in the *phyC* mutant throughout a diurnal cycle (Fig 2F). In summary, the transcriptional profiles of *FT1*, *VRN1, VRN2, CO1*, and *CO2* are similar between *elf3* and *elf3/phyC* mutants consistent with *ELF3* acting downstream from *PHYC* in the photoperiod flowering pathway.

**Fig 2.**
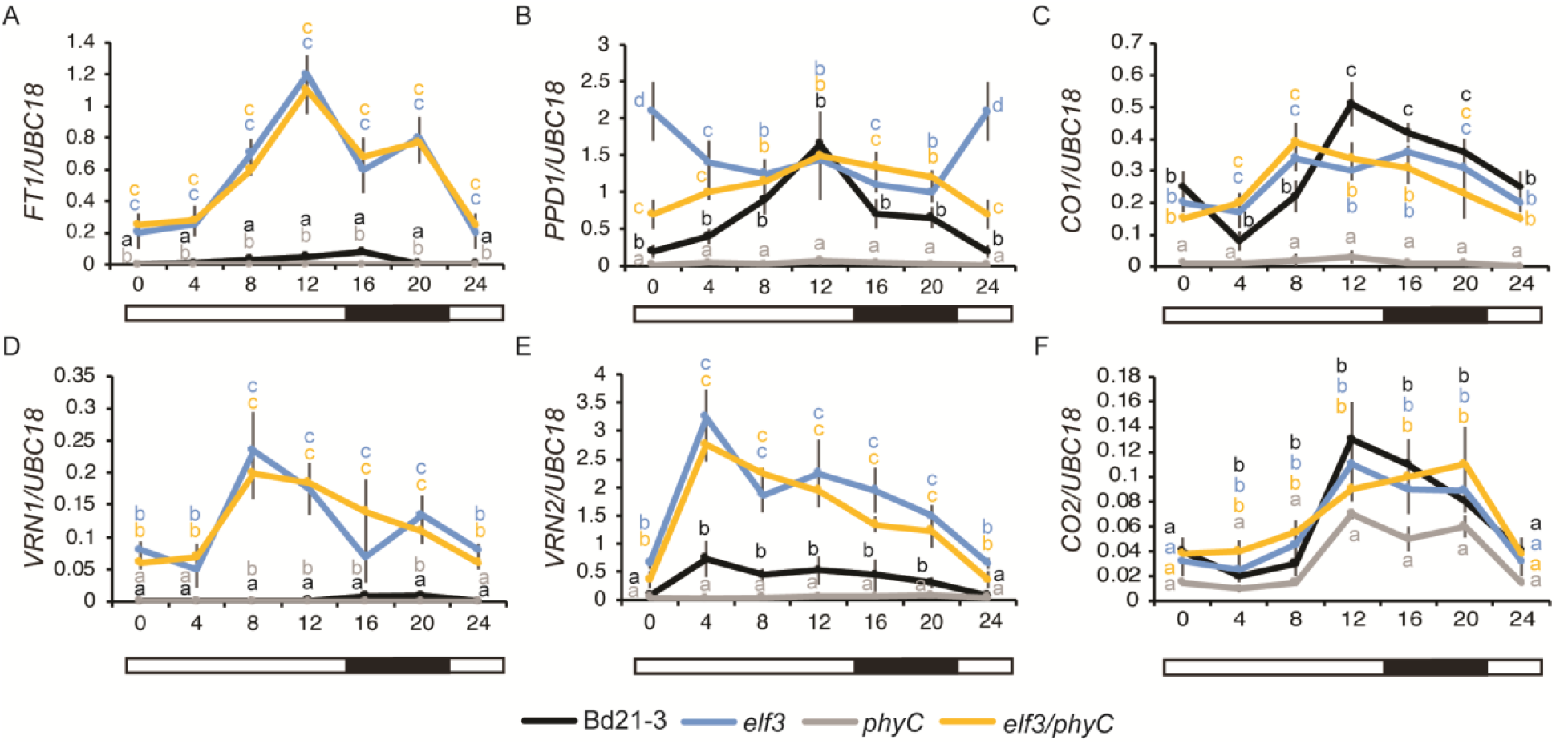
Effect of loss of function mutations in *ELF3* and *PHYC* on the transcriptional profiles of six flowering time genes in 16h long days. Normalized expression of (**A**) *FT1*, (**B**) *PPD1*, (**C**) *CO1*, (**D**) *VRN1*, (**E**) *VRN2*, and (**F**) *CO2* during a 24h diurnal cycle in Bd21-3 (black line), *elf3* (blue lines), *phyC* (gray lines) and *elf3/phyC* double mutant (orange line). Plants were grown in LDs until the fourth-leaf stage was reached (Zadoks=14) at which point the newly expanded fourth leaf was harvested at zt0, zt4, zt8, zt12, zt16, and zt20. Note the zt0 value and zt24 value are the same. The average of four biological replicates is shown (three leaves per replicate). Error bars represent the standard error of the mean. Data were normalized using *UBC18* as done in [49].

There is a more complex interaction between *PHYC* and *ELF3* on the transcriptional profile of *PPD1*. In wild-type, the expression levels of *PPD1* peak at zt12 with the lowest expression level at dawn and during the evening consistent with previous reports of *PPD1* expression patterns in *B. distachyon* [29,30] (Fig 2B). In both the *elf3* and *elf3/phyC* mutants, we observed increased *PPD1* expression relative to wild-type at dawn and during the evening with expression levels similar to wild-type at zt12. Interestingly, the increased expression of *PPD1* observed at dawn and during the night in the *elf3/phyC* background, was significantly lower than the single *elf3* mutant suggesting *PHYC* may impact *PPD1* expression via additional genes beyond *ELF3*. In contrast, *PPD1* expression levels were reduced in the *phyC* mutant relative to wild-type, *elf3*, and *elf3/phyC* mutant plants throughout a diurnal cycle, consistent with the reduced *FT1* expression and delayed flowering phenotype of the *phyC* mutant.

### Identification and mapping of a *ppd1* mutant in *B. distachyon*

To determine the role of *PPD1* in flowering in *B. distachyon* a whole genome sequenced sodium azide mutant line NaN610 with a predicted high effect mutation impacting a splice acceptor donor site in *PPD1* (BdiBd21-3.1G0218200) was ordered from the Joint Genome Institute (JGI) ([66]**; https://phytozome-next.jgi.doe.gov/jbrowse/**). A quarter of the NaN610 M3 seeds received were segregating for an extremely delayed flowering phenotype (Fig 3B-D).

**Fig 3.**
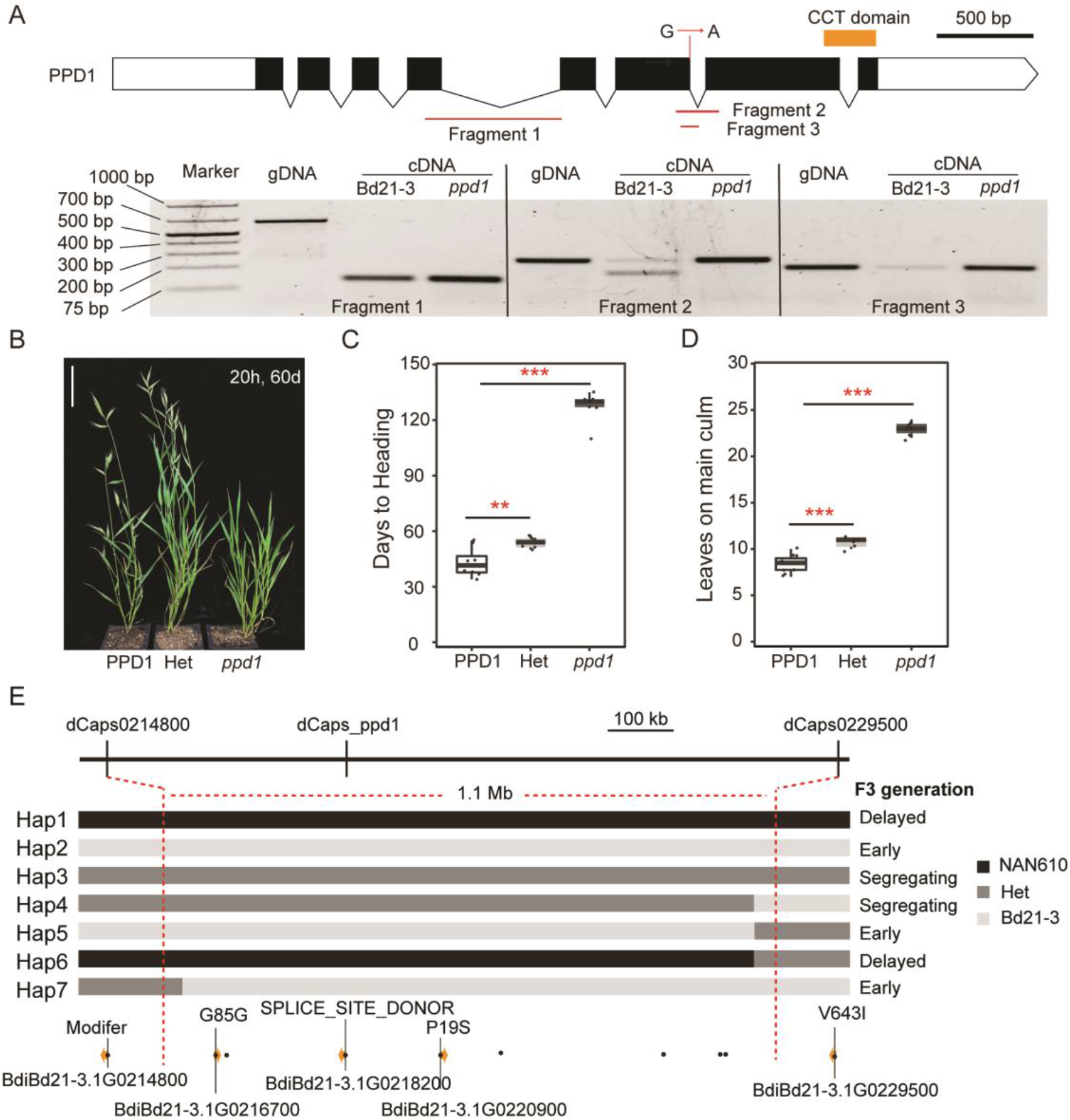
Identification of a *ppd1* mutant from an NaN line. (**A**) Gene structure of *PPD1* showing the location of the nucleotide change of the sodium azide-induced mutation, orange bar indicates the region that encodes the CCT domain. Below the gene structure diagram is a gel image of the reverse transcription polymerase chain reaction (30 cycles of amplification) showing PCR products of *PPD1* cDNA in Bd21-3 and *ppd1* mutant plants. The location of primers used in each reaction are shown in the diagram above the gel image. (**B**) Representative photo of Bd21-3, heterozygous, and homozygous *ppd1* plants grown in a 20h LD. Picture was taken after 60d after germination in 20h LD, bar=5cm. (**C** and **D**) Flowering time measured as days to heading (**C**) and the number of leaves on the parent culm at time of heading (**D**), ** indicate statistical differences (p < 0.01), *** indicate statistical differences (p < 0.001) by Student’s t-test. (**E**) Fine mapping of *ppd1* in a population of 380 BC1F2 individuals. Individuals with seven different haplotypes were identified by three dCAPS markers and flowering times of each haplotype were determined in the F3 generation. Dark, grey, and light grey rectangles represent NAN610, heterozygous, and Bd21-3 genotypes, respectively. Variants around the *PPD1* locus from NAN610 line are shown with black dots, and yellow arrow indicates coding genes within the mapped interval with the specific effect on the coding region indicated.

Due to the high mutant load of the publicly available sequenced NaN mutant lines, we validated through mapping that the delayed flowering phenotype is associated with *PPD1* (Fig 3E, F). We backcrossed NaN610 with Bd21-3 and confirmed a quarter of the plants in the BC1F2 population (n=380) were delayed flowering, demonstrating the recessive nature of the mutant. Three Derived Cleaved Amplified Polymorphic Sequences (dCAPs) markers closely linked with *PPD1* were developed based on the variant’s information for the NAN610 line, with one of the dCAPs primers located within the *PPD1* locus itself (Fig 3E; Table S1). This approach allowed us to map the causative lesion to within a 1Mb interval (13.1Mb-14.2Mb) on the top arm of chromosome 1 and indicates the delayed flowering phenotype is tightly linked with *PPD1* (Fig 3E).

To confirm that the predicted splice site mutation does in fact impact the splicing of *PPD1*, we sequenced the mRNA products of the *ppd1* NaN610 mutant line and Bd21-3 (Fig 3A). We found that the splice site mutation resulted in the mis-splicing of the sixth intron, generating a reading frame shift resulting in a truncated protein lacking the conserved CCT domain (Fig 3A). Taken together, the extreme delayed flowering of the *B. distachyon ppd1* mutant is consistent with the *ppd1* null mutants described in wheat, which take >120 days to head under inductive LD conditions [16,43], demonstrating *PPD1* is required for LD flowering broadly within temperate grasses.

### Genetic interactions between *ELF3* and *PPD1* under long and short days

We and others have shown that *PPD1 or PRR37* expression is induced in an *elf3* mutant background in *B. distachyon*, rice and wheat (Fig 2B; [29,30,40,68]). Moreover, a CHIPseq analysis of ELF3 demonstrated that *PPD1* or *PRR37* is directly bound by ELF3 in a time of day responsive manner [29,40]. Thus, ELF3 acts as a direct transcriptional repressor of *PPD1 or PRR37* but the extent to which this explains the rapid flowering in the *elf3* mutant has not been tested. Therefore, we generated an *elf3/ppd1* double mutant to explore the genetic interactions between these two genes have under a highly inductive 20h LD, inductive 16h LD and non-inductive 8h SD (Fig 4).

**Fig 4.**
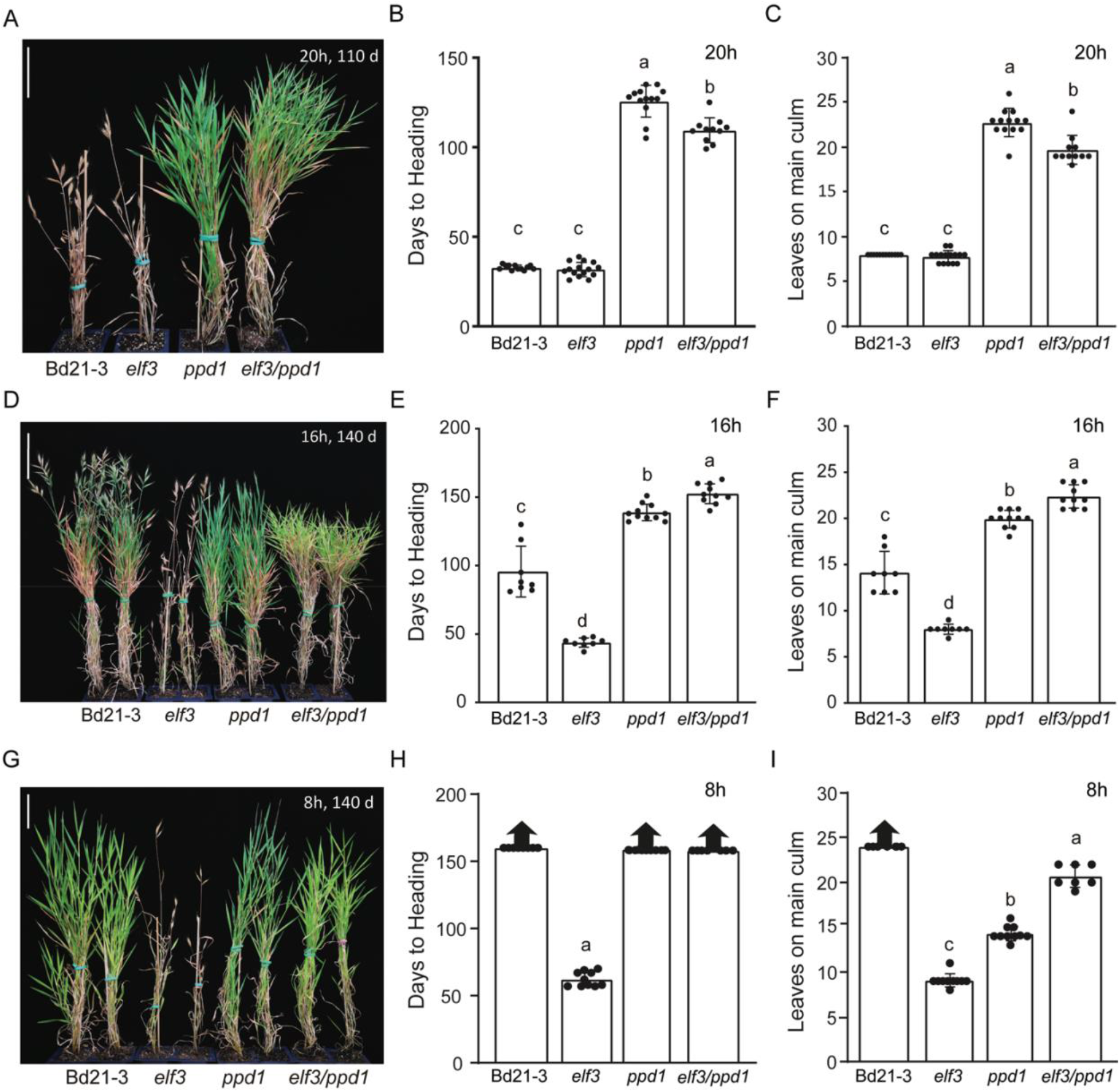
Genetic interactions between the delayed flowering *ppd1* mutant and the rapid flowering *elf3* mutant. Representative photo of Bd21-3 wild-type, rapid flowering *elf3* mutant, delayed flowering *ppd1* mutant and delayed flowering *elf3/ppd1* double mutant plants grown in a 20h photoperiod (**A**), 16h photoperiod (**D**), and 8h photoperiod (**G**). Picture was taken after 110d, for the 20h LD (**A**) and 140d after germination for the 16h LD (**D**), and 8h SD. Scale bar=5cm. (**B**, **E**, **H**) Flowering times under 20h (**B**), 16h (**E**), 8h (**G**) measured as days to heading of Bd21-3, *elf3, ppd1*, and *elf3/ppd1*. Flowering times under 20h (**C**), 16h (**F**), and 8h (**I**) measured as the number of leaves on the parent culm at time of heading. Bars represent the average of 8 plants ± SD. Arrows above bars indicate that none of the plants flowered at the end of the experiment (150d). Letters (a, b, c, d) indicate statistical differences (p < 0.05) according to a Tukey’s HSD test used to perform multiple comparisons

Under all photoperiods the *elf3/ppd1* double mutant flowered significantly later than the *elf3* single mutant plants (Fig 4). Interestingly, under 20h LD *elf3/ppd1* double mutant plants flowered significantly earlier than *ppd1* by 16.2 days forming 3.0 fewer leaves whereas under both 16h or 8h days *elf3/ppd1* mutant plants flowered significantly later than *ppd1* by 13.7 days with 2.5 more leaves in 16h days and with 6.4 more leaves in 8h days (Fig 4D-I). Also, surprisingly while *ppd1* mutant plants are delayed in flowering compared to wild-type under 20h and 16h LD, under 8h SD when using leaf number as a metric for flowering indicates that *ppd1* mutant plants transition to flower earlier under 8h SD than wild-type and *elf3/ppd1* double mutant plants (Fig 4I). Even though *ppd1* mutant plants stop producing leaves these plants still fail to head after 160 days of growth when the experiment was terminated (Fig 4H). Additionally, *elf3/ppd1* double plants while heading later than *ppd1* single mutant plants still transition to flower earlier than wild-type producing fewer leaves than wild-type in SD but still failing to head (Fig 4H-I). It is also worth noting that *elf3/ppd1* double mutant plants are still able to respond to different photoperiods, with longer days resulting in significantly earlier flowering plants than under shorter days (Fig 4B, E, H). Taken together, these results indicate that there are strong genetic interactions between *ELF3* and *PPD1* under different photoperiods, that *PPD1* is a key downstream gene of *ELF3*, and that there is a residual photoperiodic response that is independent of these two genes.

### Effect of mutations in *ELF3* and *PPD1* on the transcriptional profiles of flowering time genes

To understand how *ELF3* and *PPD1* affect flowering at a molecular level, we measured the mRNA levels of *FT1*, *VRN1, PPD1, VRN2, CO1*, and *CO2* in the *elf3* and *ppd1* single mutants and the *elf3/ppd1* double mutant across a diurnal cycle in 16h LD (Fig 5). As before, *FT1* and *VRN1* expression levels were elevated in the *elf3* mutant background however, in the *elf3/ppd1* double mutant expression of these genes remained low and resembled the expression profile of *ppd1* single mutants (Fig 5A, D). The low expression levels of *FT1* and *VRN1* in *ppd1* and *ppd1/elf3* mutant plants is consistent with the delayed flowering phenotype for both of these mutants in 16h long days. The *VRN2* expression profile was similar between wild-type and *ppd1* mutant plants with low expression levels at dawn and increased expression throughout the light cycle before expression levels dropping in the dark (Fig 5E). Interestingly, *VRN2* expression levels are similarly elevated throughout the day in *elf3* and *elf3/ppd1* mutant plants compared to wild-type (Fig 5E).

**Fig 5.**
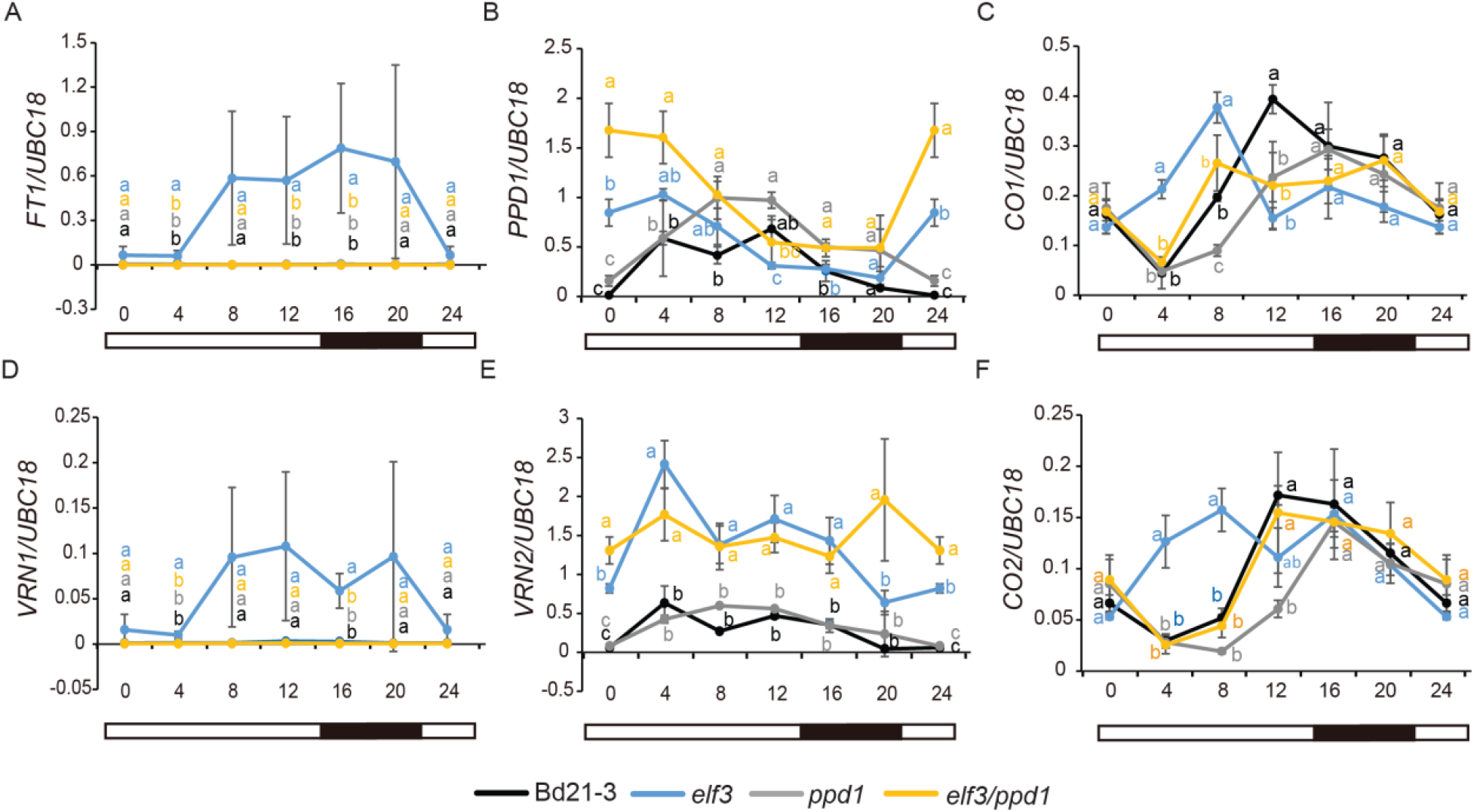
Effect of loss of function mutations in *ELF3*, and *PPD1* on the transcriptional profiles of six flowering time genes in 16h long days. The fourth newly expanded leaves were harvested every 4h over a 24-hour period, three biological replicates (two leaves per replicate) were harvested at each time point for each genotype. Diurnal expression of *FT1* (**A**), *PPD1* (**B**), *CO1* (**C**), *VRN1* (**D**), *VRN2* (**E**), and *CO2* (**F**) were detected in Bd21-3 (black line), *elf3* (blue lines), *ppd1* (grey lines) and *elf3/ppd1* double (orange line), respectively. Bars represent the average of three biological replicates ± SD. Letters (a, b, c, d) indicate statistical differences (p < 0.05) according to a Tukey’s HSD test used to perform multiple comparisons, letter color corresponds to the four different genotypes.

Consistent with the expression patterns of *PPD1* in wild-type and *elf3* single mutant shown in Fig. 2, the expression levels of *PPD1* peak at zt12 in wild-type and in the *elf3* mutant there is increased *PPD1* expression relative to wild-type at dawn and during the evening (Fig 5B). *PPD1* expression levels in the *ppd1* mutant should be interpreted with caution because we do not know the effect of the splice site mutation on the mRNA stability. Significantly higher levels of *PPD1* expressions were observed in *ppd1* relative to wild-type at ZT8 and ZT16, and in *elf3/ppd1* relative to *elf3* at dawn. But the expression patterns of *PPD1* were more similar between *ppd1* and wild-type and between *elf3/ppd1* and *elf3* than across these comparisons (Fig 5B).

*CO1* and *CO2* expression both have peak expression in wild-type at zt12 with expression dampening in the evening consistent with previous reports [29,58].

Interestingly, the expression levels of *CO1* and *CO2* were elevated between zt4-8 in the *elf3* mutant compared with wild-type however from zt 12-20 expression levels were lower than wild-type with a profile similar to *elf3/ppd1* mutant plants. In contrast the expression levels of *CO1* and *CO2* were the lowest in the *ppd1* plants compared with all the other lines at zt8. In the *elf3/ppd1* mutant plants *CO1* and *CO2* expression were more similar to *ppd1* in the morning and to *elf3* in the evening and night (Fig 5C, F). These results indicate complex interactions between PPD1 and ELF3 in the regulation of *CO1* and *CO2*.

### Constitutive expression of *ELF3* results in delayed flowering and a reduction of *PPD1, FT1*, and *VRN1* expression levels

In our previous study, we showed that overexpression of *ELF3* in the *elf3* mutant background results in delayed flowering taking nearly 5 months to flower ([30], Fig6 A). However, this was done in the T0 generation, so here we evaluated the flowering time and expression of downstream flowering time genes in the T1 generation. We grew four *UBI::ELF3/elf3* transgenic lines alongside Bd21-3 and *elf3* in a 16h photoperiod, and harvested the newly expanded fourth leaf at zt4. This time point was chosen because expression of several critical genes such as *CCA1*, *TOC1, LUX, PPD1, VRN2, CO1*, and *CO2* were significantly different at dawn in the *elf3* single mutant compared with wild-type [29,30];Fig 2 and Fig 5). We first confirmed that all the *UBI::ELF3/elf3* transgenic lines had higher ELF3 levels, and found indeed there is a significant increase of expression of *ELF3* in the transgenic plants (Fig 6B). To understand how *UBI::ELF3/elf3* affects flowering time, we detected the expression of *FT1*, *VRN1, PPD1, VRN2, CO1*, and *CO2* in wild-type, *elf3* and, *UBI::ELF3/elf3*. Consistent with the delayed flowering time, *FT1* and *VRN1* expression levels of *UBI::ELF3/elf3* were reduced compared to elevated levels in *elf3* single mutant plants (Fig 6 C and F). Also, the expression of *PPD1, VRN2, CO1*, and *CO2* were decreased in *UBI::ELF3/elf3*, indicating ELF3 is playing a repressive role broadly in regulating CCT domain containing genes responding to photoperiodic flowering.

**Fig 6.**
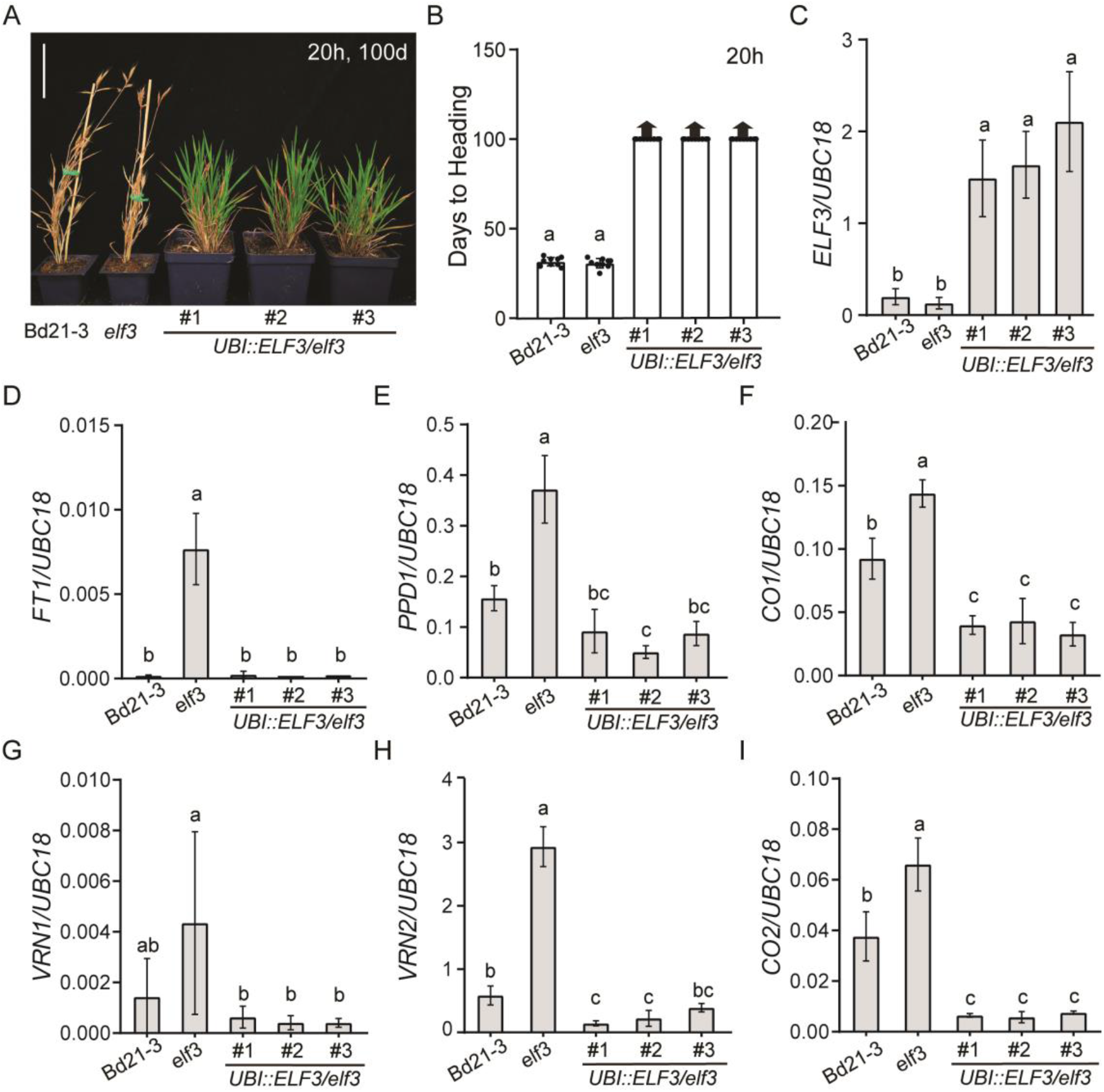
Overexpression of *ELF3* in the *elf3* mutant background delays flowering. (**A**) Representative photo of Bd21-3 wild-type, rapid flowering *elf3* mutant and three independent transgenic plants of *UBI::ELF3* in the *elf3* background grown in a 20h photoperiod. The fourth newly expanded leaves were harvested at zt4 in 16h. Picture was taken 100d after germination. Bar = 5 cm. (**B**-**I**), Normalized expression of *ELF3* (**C**), *FT1* (**D**),*PPD1* (**E**), *CO1* (**F**), *VRN1* (**G**), *VRN2* (**H**) and *CO2* (**I**) were detected in Bd21-3, *elf3* single mutant and four *UBI::ELF3/elf3* transgenic lines. Bars represent the average of four biological replicates ± SD.

## DISCUSSION

### The phytochromes *PHYC/PHYB* and *ELF3* connection

*B. distachyon* has an obligate requirement for LD to flower [8,49,67]. Previous studies have shown the important roles that both PHYC and ELF3 play in photoperiodic flowering in *B. distachyon* [29,30,58]. Specifically, mutations in *phyC* result in extremely delayed flowering whereas loss-of-function mutations in *elf3* results in rapid flowering in either LD or SD [30,58]. Furthermore, *phyC* mutants resemble plants grown in SD both morphologically and at the transcriptomic level regardless of day-length whereas *elf3* mutant plants resemble plants grown in LD both morphologically and at the transcriptomic level regardless of day-length [29,30,58]. Thus, we were interested in exploring the genetic relationships between *PHYC* and *ELF3*. The extreme delayed flowering phenotype observed in *phyC* mutant plants is mediated by *ELF3* because *phyC/elf3* double mutant plants flower rapidly in LD and SD similar to *elf3* mutant plants. Similar genetic interactions between *phyB* and *elf3* were also found in wheat in the companion study [68], suggesting these interactions are likely to be conserved broadly in temperate grasses. Loss-of-function mutations in *phyB* in wheat also result in delayed flowering similar to *phyC* [59]. At present, no null *phyB* alleles have been reported in *B. distachyon;* however, PHYB is able to heterodimerize with PHYC in *B. distachyon* and wheat [29,57] and both *phyB* [59] and *phyC* [57] are extremely late flowering in wheat suggesting that both PHYs are likely required for photoperiodic flowering in the temperate grasses.

Phytochrome regulation of ELF3 at the post-translational level rather than at the transcriptional level is likely to be the critical interaction impacting flowering. In *A. thaliana*, *B. distachyon*, and wheat, *phyB/phyC* mutants do not impact the circadian oscillation of *ELF3* mRNA levels [29,59,69]. However, in all three species PHYB and PHYC have been shown to interact with the ELF3 protein, but the stability of the ELF3 protein upon exposure to light differs between *A. thaliana* and temperate grasses [29,68,70,71]. Specifically, in *A. thaliana* PHYB contributes to the stability of the ELF3 protein during the light leading to ELF3 accumulation at the end of the day whereas in temperate grasses ELF3 protein accumulates during the night and is rapidly degraded/modified upon light exposure [29,70,72,73]. We hypothesize that the rapid degradation and or modification of ELF3 by light is likely PHY mediated in temperate grasses as shown in rice [40].

The differences in how phytochromes impact the stability of the ELF3 protein might explain the contrasting flowering phenotypes of the *phyB/phyC* mutants between *A. thaliana* and the temperate grasses. In *A. thaliana, phyB* mutants flower more rapidly than wild type in either LD or SD and *phyC* mutants flower earlier under SD [60], whereas in temperate grasses *phyB* or *phyC* mutants are extremely delayed in flowering [57–59]. However, ELF3 acts as a flowering repressor in both *A. thaliana* and temperate grasses [30,32]. In *A. thaliana* PHYB stabilizes the ELF3 protein; therefore, in *phyB* mutants, ELF3 is no longer stable leading to rapid flowering. In contrast, in temperate grasses, in the absence of *phyB* or *phyC* the ELF3 protein is more stable leading to delayed flowering.

Interestingly, overexpression of ELF3 results in extremely delayed flowering in *B. distachyon* [29,30] (Fig 6). Given that the regulation of ELF3 appears to occur at the protein level, one might not expect that overexpression would cause such a strong flowering delay. However, if the ELF3 protein is expressed at a high level such that the degradation machinery is unable to degrade most of the ELF3 protein during LD, then a strong flowering delay might occur. In support of this idea, the delayed flowering of overexpression of ELF3 is mitigated when plants are grown under constant light versus normal 16h LD conditions [29]. It is worth noting that although overexpression of ELF3 generally leads to delayed flowering in different plant species, there is considerable variation in the magnitude of this delayed flowering [29,63,68] (Fig 6).

Similar genetic interactions between *ELF3* and *PHYB* have also been observed in rice which is a SD-flowering plant that has two rice-specific *ELF3* paralogs [73]. Mutations in either paralog results in delayed flowering in SD or LD in contrast to the rapid flowering observed in *elf3* mutations in temperate grasses [40,74,75]. Also in contrast to the situation in temperate grasses, *phyB* mutants flower more rapidly than wild type in rice [76]. Despite the flowering differences of the *elf3* and *phyB* mutants between rice and temperate grasses, in either case the light signal perceived by the phytochromes is ELF3 mediated [75]. Specifically, in both rice and temperate grasses, in *phyB/elf3* or *phyC/elf3* mutant plants, *elf3* is epistatic to *phyC*. Moreover, PHYB and ELF3 proteins interact impacting the modification of ELF3 by light [40]. The opposite roles that phytochromes and *elf3* have on flowering in rice and temperate grasses is likely in part due to the reverse role that the downstream *PPD1/PRR37* gene has on flowering because *PPD1* is a promoter of flowering in LD temperate grasses but is a repressor of flowering in SD grasses such as rice [11,16,77,78] (Fig 3).

### The *ELF3* and *PPD1* connection

The extremely delayed flowering of *B. distachyon ppd1* mutant plants under LD is similar to the extremely delayed heading of *ppd1* mutants in wheat [16]. However, a previous study in *B. distachyon* using a CRISPR induced *ppd1* mutant allele which has a 1bp deletion in the sixth exon of *PPD1* has a milder delayed flowering phenotype with plants taking around 40 days to flower under 20h days, whereas the mutant *ppd1* plants presented here flower around 120 days in 20h days [29]; Fig 3, 4). In both studies, wild-type Bd21-3 plants flower on average between 25-30 days in 20h of light consistent with previous reports in *B. distachyon* [49,79]. The differences in flowering time between the two *B. distachyon ppd1* mutant alleles suggests that the CRISPR induced *ppd1* allele is a weaker hypomorphic allele than the *ppd1* mutant allele characterized in this study. This is further supported by the fact that the *ppd1* allele described here has a similar extremely delayed flowering phenotype as the null *ppd1* wheat allele [16].

Interestingly, *B. distachyon ppd1* mutant plants flower in SD (8 h day-length) in contrast to wild type which cannot flower in SD. Indeed, all 56 different *B. distachyon* accessions evaluated in SD remained vegetative after >200 days of growth [49,67,79]. In previous studies, we showed that wild-type *B. distachyon* has an obligate requirement for LD of at least 12 hours to flower [49,80]. The inability of *B. distachyon* to flower in SD has been evaluated not only by the appearance of visible flowers, but also by dissecting meristems to determine if floral primordia have formed. We note the need for microscopic examination of dissected meristems for the presence of floral primordia to determine, from a developmental perspective, if flowering has occurred because, in the agronomic literature on cereals, flowering is often evaluated as days to heading, and there are many examples of genotypes that are “non-flowering” in SD that have actually formed floral primordia, but further floral development is arrested until the plants are in LD. For example, photosensitive wheat and *ppd1* mutant plants fail to head under 8h SD [16, 68]; however, dissection of meristems reveal that photosensitive wheat and *ppd1* actually transition to flowering by leaf 7-8 in SD [16], suggesting contrasting effects of *PPD1* in SD in *B. distachyon* and wheat. However, this contrasting effect may be wheat genotype dependent: spring wheats exhibit a microscopic flowering transition in SD whereas certain winter wheat do not (e.g. [81]), but perhaps a *pdd1* mutation in winter wheat would result in a microscopic flowering transition in SD (i.e., perhaps in winter wheat there would not be contrasting effects with *B. distachyon* (winter accession) of a *ppd1* lesion in SD).

The *ppd1/elf3* double mutant is delayed in flowering relative to *elf3* mutant plants indicating that *PPD1* is downstream of ELF3 for photoperiodic flowering. This is also consistent with the elevated *PPD1* expression levels observed at dawn and dusk in the *elf3* mutant relative to wild-type in temperate grasses [30,73] (Fig 5). Indeed, ELF3 binds to the *PPD1/PRR37* promoter in *B. distachyon*, wheat, and rice indicating ELF3 is a direct transcriptional repressor of *PPD1/PRR37* in grasses [29,40,68]. ELF3 does not have any known DNA binding activity and thus, the direct repression is likely to be due to ELF3’s interaction with the LUX transcription factor which, from studies in *A. thaliana*, recognizes GATWCG motifs that are also found in the *PPD1* promoter in grasses [37,38,68]. Interestingly, photoperiod insensitivity in wheat is associated with deletions in the *PPD1* promoter, which remove the LUX binding site and results in elevated *PPD1* expression at dawn similar to the *PPD1* expression dynamics observed in *elf3* and *lux* mutant plants [10,12,13, 39,82]. In the companion paper, ChIP-PCR experiments show ELF3 enrichment around the LUX binding site, which is present within the region deleted in photoperiod-insensitive wheats. These results demonstrate that removal of the ELF3 binding site leads to elevated expression in *PPD1* and accelerated heading under SD in many photoperiod-insensitive wheats [68].

The characterization of *elf3/ppd1* mutant plants under different photoperiods reveal complex interactions between the two genes and their downstream targets depending on the environment. For example, *elf3/ppd1* mutant plants are earlier flowering than *ppd1* mutant plants under 20h days but, surprisingly, are later flowering under 16h and 8h days. This is in contrast with *elf3/ppd1* mutants in wheat, which head earlier than *ppd1* under either 16h-LD or 8h-SD photoperiods indicating that *ELF3* can delay heading independently of PPD1 in both photoperiods [68]. Thus, there are differences between *B. distachyon* and wheat in the effects of *ELF3* on heading in the absence of *PPD1*. We speculate that these differences may be related to the different interactions observed between *CO1* and other flowering genes (e.g. PHY) in these two species. For example, in wheat *phyC* and *ppd1* mutants, *CO1* expression levels are elevated compared to wild-type, whereas in *B. distachyon CO1* expression is reduced in both mutants [57, 58, 68].

## MATERIALS AND METHODS

### Plant Materials and Growth Conditions

The rapid flowering mutant *elf3* and four *UBI::ELF3/elf3* transgenic lines were previously characterized and generated in [30] and the delayed flowering *phyC* mutant was characterized in [58]. All the mutants used for phenotyping and expression in this study were backcrossed at least twice with the wild-type Bd21-3 accession. Seeds were imbibed in the dark at 5°C for three days before placing seeds into soil. Three photoperiods 8h-SD (8h light/16h dark), 16h-LD (16h light/ 8h dark), and 20h-LD (20h light/ 4h dark) were used, and growth conditions (temperature, light) for data in figures 3-4 were described in [49] whereas the growth conditions for Figure 1-2 are described in [16]. Flowering time was estimated by measuring days to heading and leaves on the main culm at time of heading. Recording of days to heading was done as the days from seed germination to the first emergence of the spike.

### Generation of *elf3/phyC* and *elf3/ppd1* double-mutant lines

Epistasis analysis between *phyC, ppd1*, and *elf3* was studied by generating an *elf3/phyC* and *elf3/ppd1* double mutant. *phyC* mutants were crossed with *elf3* and *elf3/phyC* homozygous double mutant plants were selected in a segregating F2 population by genotyping using primers in Table S1. Similarly, *ppd1* was crossed with *elf3*, and *elf3/ppd1* homozygous double mutant individuals were selected by genotyping using primers in Table S1 in the segregating F2 population. Flowering time of *elf3/ppd1* double mutant were estimated by growing with Bd21-3, *elf3, ppd1* side by side in 8h SD, 16h LD, and 20h LD whereas *elf3/phyC* double mutant plants were grown in 8h SD and 16h LD.

### RNA Extraction and qPCR

The method for RNA extraction, cDNA synthesis and quantitative PCR (qPCR) is described in [49]. Primers used for gene expression analyses are listed in Table S1.

### Statistical Analyses

For more than two genotypes comparison, Multiple comparisons were performed by using *agricolae* package in R [83]. Statistically significant differences among different genotypes were calculated by using one-way analysis of variance (ANOVA) followed by a Tukey’s HSD test. Student’s t-test was used for analyzing the difference between two genotypes, significant if P< 0.05.

## ACKNOWLEDGMENTS

Thanks to Frédéric Bouché for fruitful discussions about photoperiod sensing in *B. distachyon*. This work was funded in part by the US National Science Foundation (Award IOS-1755224) and by the Great Lakes Bioenergy Research Center, US Department of Energy, Office of Science, Office of Biological and Environmental Research (Award No. DE-SC0018409). The work from JV, RS, and MS (proposal: 10.46936/10.25585/60001041) conducted by the U.S. Department of Energy Joint Genome Institute (https://ror.org/04xm1d337), a DOE Office of Science User Facility, is supported by the Office of Science of the U.S. Department of Energy operated under Contract No. DE-AC02-05CH11231. DPW was a Howard Hughes Medical Institute (HHMI) Fellow of the Life Sciences Research Foundation which supported the work while in Jorge Dubcovsky’s lab. JD was funded by HHMI. DL is funded by USDA-ARS CRIS Project 2030-21430-014D.

## Supporting Information Captions

**Table S1.**
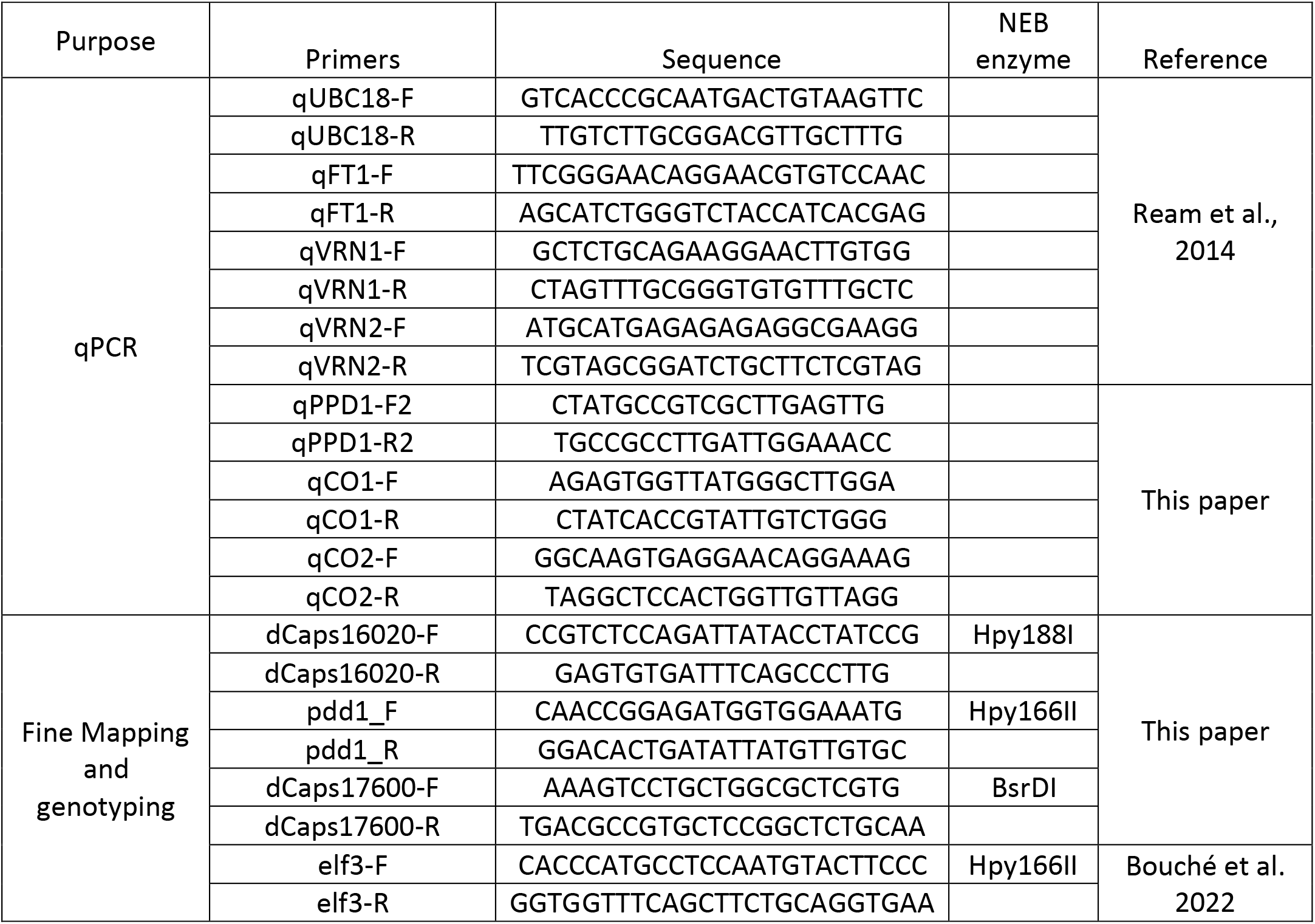
Primers used in this study

## REFERENCES

1. Andrés F, Coupland G. The genetic basis of flowering responses to seasonal cues. Nat Rev Genet. 2012;13: 627–639. doi:10.1038/nrg3291

2. Fjellheim S, Boden S, Trevaskis B. The role of seasonal flowering responses in adaptation of grasses to temperate climates. Front Plant Sci. 2014;5: 431. doi:10.3389/fpls.2014.00431

3. Bouché F, Woods DP, Amasino RM. Winter Memory throughout the Plant Kingdom: Different Paths to Flowering. Plant Physiol. 2017;173: 27–35. doi:10.1104/pp.16.01322

4. Gaudinier A, Blackman BK. Evolutionary processes from the perspective of flowering time diversity. New Phytol. 2020;225: 1883–1898. doi:10.1111/nph.16205

5. Bendix C, Marshall CM, Harmon FG. Circadian Clock Genes Universally Control Key Agricultural Traits. Molecular Plant. 2015;8: 1135–1152. doi:10.1016/j.molp.2015.03.003

6. Song YH, Shim JS, Kinmonth-Schultz HA, Imaizumi T. Photoperiodic Flowering: Time Measurement Mechanisms in Leaves. Annu Rev Plant Biol. 2015;66: 441–464. doi:10.1146/annurev-arplant-043014-115555

7. Woods DP, Amasino RM. Dissecting the Control of Flowering Time in Grasses Using *Brachypodium distachyon*. Genetics and Genomics of Brachypodium. Cham: Springer International Publishing; 2015. pp. 259–273. doi:10.1007/7397_2015_10

8. Raissig MT, Woods DP. The wild grass *Brachypodium distachyon* as a developmental model system. Curr Top Dev Biol. 2022; 147: 33–71. doi:10.1016/bs.ctdb.2021.12.012

9. Hasterok R, Catalán P, Hazen SP, Roulin AC, Vogel JP, Wang K, et al. Brachypodium: 20 years as a grass biology model system; the way forward? Trends in Plant Science. 2022;27: 1002–1016. doi:10.1016/j.tplants.2022.04.008

10. Beales J, Turner A, Griffiths S, Snape JW, Laurie DA. A Pseudo-Response Regulator is misexpressed in the photoperiod insensitive *Ppd-D1a* mutant of wheat (*Triticum aestivum L.*). Theor Appl Genet. 2007;115: 721–733. doi:10.1007/s00122-007-0603-4

11. Turner A, Beales J, Faure S, Dunford RP, Laurie DA. The pseudo-response regulator *Ppd-H1* provides adaptation to photoperiod in barley. Science. 2005;310: 1031–1034. doi:10.1126/science.1117619

12. Campoli C, Shtaya M, Davis SJ, Korff von M. Expression conservation within the circadian clock of a monocot: natural variation at barley *Ppd-H1* affects circadian expression of flowering time genes, but not clock orthologs. BMC Plant Biol. 2012;12: 1–1. doi:10.1186/1471-2229-12-97

13. Wilhelm EP, Turner AS, Laurie DA. Photoperiod insensitive *Ppd-A1a* mutations in tetraploid wheat (*Triticum durum Desf*.). Theor Appl Genet. 2008;118: 285–294. doi:10.1007/s00122-008-0898-9

14. Shaw LM, Turner AS, Herry L, Griffiths S, Laurie DA. Mutant Alleles of *Photoperiod-1* in Wheat (*Triticum aestivum L*.) That Confer a Late Flowering Phenotype in Long Days. PLoS ONE. 2013;8: e79459. doi:10.1371/journal.pone.0079459.s003

15. Seki M, Chono M, Matsunaka H, Fujita M, Oda S, Kubo K, et al. Distribution of photoperiod-insensitive alleles *Ppd-B1a* and *Ppd-D1a* and their effect on heading time in Japanese wheat cultivars. Breed Sci. 2011;61: 405–412. doi:10.1270/jsbbs.61.405

16. Shaw LM, Li C, Woods DP, Alvarez MA, Lin H, Lau MY, et al. Epistatic interactions between *PHOTOPERIOD1*, *CONSTANS1* and *CONSTANS2* modulate the photoperiodic response in wheat. PLoS Genet. 2020;16: e1008812–28. doi:10.1371/journal.pgen.1008812

17. Woods DP, Bednarek R, Bouché F, Gordon SP, Vogel JP, Garvin DF, et al. Genetic Architecture of Flowering-Time Variation in *Brachypodium distachyon*. Plant Physiol. 2017;173: 269–279. doi:10.1104/pp.16.01178

18. Bettgenhaeuser J, Corke FMK, Opanowicz M, Green P, Hernández-Pinzón I, Doonan JH, et al. Natural Variation in Brachypodium Links Vernalization and Flowering Time Loci as Major Flowering Determinants. Plant Physiol. 2017;173: 256–268. doi:10.1104/pp.16.00813

19. Gordon SP, Contreras-Moreira B, Woods DP, Marais Des DL, Burgess D, Shu S, et al. Extensive gene content variation in the *Brachypodium distachyon* pan-genome correlates with population structure. Nature Communications. 2017;8: 1–13. doi:10.1038/s41467-017-02292-8

20. Tyler L, Lee SJ, Young ND, DeIulio GA, Benavente E, Reagon M, et al. Population Structure in the Model Grass Is Highly Correlated with Flowering Differences across Broad Geographic Areas. The Plant Genome. 2016;9: 0–20. doi:10.3835/plantgenome2015.08.0074

21. Wilson P, Streich J, Borevitz J. Genomic Diversity and Climate Adaptation in Brachypodium. Genetics and Genomics of Brachypodium. Cham: Springer International Publishing; 2015. pp. 107–127.

22. Zikhali M, Wingen LU, Griffiths S. Delimitation of the *Earliness per se D1* (*Eps-D1*) flowering gene to a subtelomeric chromosomal deletion in bread wheat (*Triticum aestivum*). Journal of Experimental Botany. 2015;67: 287–299. doi:10.1093/jxb/erv458

23. Alvarez MA, Tranquilli G, Lewis S, Kippes N, Dubcovsky J. Genetic and physical mapping of the earliness per se locus *Eps-Am1* in *Triticum monococcum* identifies *EARLY FLOWERING 3* (*ELF3*) as a candidate gene. Funct Integr Genomics. 2016; 1–18. doi:10.1007/s10142-016-0490-3

24. Faure S, Turner AS, Gruszka D, Christodoulou V, Davis SJ, Korff von M, et al. Mutation at the circadian clock gene *EARLY MATURITY 8* adapts domesticated barley (*Hordeum vulgare*) to short growing seasons. Proc Natl Acad Sci USA. 2012;109: 8328–8333. doi: 10.1073/pnas.1120496109.

25. Zakhrabekova S, Gough SP, Braumann I, Müller AH, Lundqvist J, Ahmann K, et al. Induced mutations in circadian clock regulator Mat-a facilitated short-season adaptation and range extension in cultivated barley. Proc Natl Acad Sci USA. 2012;109: 4326–4331. doi:10.1073/pnas.1113009109/-/DCSupplemental/st01.doc

26. Matsubara K, Ogiso-Tanaka E, Hori K, Ebana K, Ando T, Yano M. Natural variation in *Hd17*, a homolog of Arabidopsis *ELF3* that is involved in rice photoperiodic flowering. Plant and Cell Physiology. 2012;53: 709–716. doi:10.1093/pcp/pcs028

27. Lundqvist, U. Eighty years of Scandinavian barley mutation genetics and breeding. In Induced Plant Mutations in the Genomics Era, Q.Y. Shu, ed. (Rome: Food and Agriculture Organization of the United Nations), 2009 pp. 39–43.

28. Hicks KA, Albertson TM, Wagner DR. *EARLY FLOWERING3* encodes a novel protein that regulates circadian clock function and flowering in Arabidopsis. Plant Cell. 2001;13: 1281–1292. doi:10.1105/tpc.13.6.1281

29. Gao M, Geng F, Klose C, Staudt A-M, Huang H, Nguyen D, et al. Phytochromes measure photoperiod in Brachypodium. bioRxiv. 2019;16: 365–53. doi:10.1101/697169

30. Bouché F, Woods DP, Linden J, Li W, Mayer KS, Amasino RM, et al. *EARLY FLOWERING 3* and Photoperiod Sensing in *Brachypodium distachyon*. Front Plant Sci. 2021;12: 769194. doi:10.3389/fpls.2021.769194

31. Huang H, Gehan MA, Huss SE, Alvarez S, Lizarraga C, Gruebbling EL, et al. Cross-species complementation reveals conserved functions for *EARLY FLOWERING 3* between monocots and dicots. Plant Direct. 2017;1: e00018–14. doi:10.1002/pld3.18

32. Huang H, Nusinow DA. Into the Evening: Complex Interactions in the Arabidopsis Circadian Clock. Trends in Genetics. 2016;32: 674–686. doi:10.1016/j.tig.2016.08.002

33. Covington MF, Panda S, Liu XL, Strayer CA, Wagner DR, Kay SA. ELF3 modulates resetting of the circadian clock in Arabidopsis. Plant Cell. 2001;13: 1305–1315. doi:10.1105/tpc.13.6.1305

34. Nusinow DA, Helfer A, Hamilton EE, King JJ, Imaizumi T, Schultz TF, et al. The ELF4–ELF3–LUX complex links the circadian clock to diurnal control of hypocotyl growth. Nature. 2011;475: 398–402. doi:10.1038/nature10182

35. Hazen SP, Schultz TF, Pruneda-Paz JL, Borevitz JO, Ecker JR, Kay SA. *LUX ARRHYTHMO* encodes a Myb domain protein essential for circadian rhythms. Proc Natl Acad Sci USA. 2005;102: 10387–10392. doi:10.1073/pnas.0503029102

36. Doyle MR, Davis SJ, Bastow RM, McWatters HG, Kozma-Bognar L, Nagy F, et al. The *ELF4* gene controls circadian rhythms and flowering time in Arabidopsis thaliana. Nature. 2002;419: 74–77. doi:10.1038/nature00954

37. Silva CS, Nayak A, Lai X, Hutin S, Hugouvieux V, Jung J-H, et al. Molecular mechanisms of Evening Complex activity in Arabidopsis. Proc Natl Acad Sci USA. 2020;117: 6901–6909. doi:10.1073/pnas.1920972117

38. Ezer D, Jung J-H, Lan H, Biswas S, Gregoire L, Box MS, et al. The evening complex coordinates environmental and endogenous signals in Arabidopsis. Nature Plants. 2017; 1–12. doi:10.1038/nplants.2017.87

39. Mizuno N, Kinoshita M, Kinoshita S, Nishida H, Fujita M, Kato K, et al. Loss-of-Function Mutations in Three Homoeologous *PHYTOCLOCK 1* Genes in Common Wheat Are Associated with the Extra-Early Flowering Phenotype. PLoS ONE. 2016;11: e0165618. doi:10.1371/journal.pone.0165618.s004

40. Andrade L, Lub Y, Cordeiro A, Costa JMF, Wigge PA, Saibo NJM, et al. The evening complex integrates photoperiod signals to control flowering in rice. Proc Natl Acad Sci USA. 2022;119:e2122582119. doi:10.1073/pnas.2122582119

41. Yan L, Fu D, Li C, Blechl A, Tranquilli G, Bonafede M, et al. The wheat and barley vernalization gene *VRN3* is an orthologue of *FT*. Proc Natl Acad Sci USA. 2006;103: 19581–19586. doi:10.1073/pnas.0607142103

42. Valverde F. CONSTANS and the evolutionary origin of photoperiodic timing of flowering. Journal of Experimental Botany. 2011;62: 2453–2463. doi:10.1093/jxb/erq449

43. Pearce S, Shaw LM, Lin H, Cotter JD, Li C, Dubcovsky J. Night-break experiments shed light on the *Photoperiod 1*-mediated flowering. Plant Physiol. 2017: pp.00361.2017–45. doi:10.1104/pp.17.00361

44. Qin Z, Bai Y, Muhammad S, Wu X, Deng P, Wu J, et al. Divergent roles of *FT-like 9* in flowering transition under different day lengths in *Brachypodium distachyon*. Nature Communications. 2019;10: 1–10. doi:10.1038/s41467-019-08785-y

45. Yan L, Loukoianov A, Tranquilli G, Helguera M, Fahima T, Dubcovsky J. Positional cloning of the wheat vernalization gene *VRN1*. Proc Natl Acad Sci USA. 2003;100: 6263–6268. doi:10.1073/pnas.0937399100

46. Li C, Dubcovsky J. Wheat FT protein regulates *VRN1* transcription through interactions with FDL2. Plant J. 2008;55: 543–554. doi:10.1111/j.1365-313X.2008.03526.x

47. Yan L, Loukoianov A, Blechl A, Tranquilli G, Ramakrishna W, San Miguel P, et al. The wheat *VRN2* gene is a flowering repressor down-regulated by vernalization. Science. 2004;303: 1640–1644. doi:10.1126/science.1094305

48. Distelfeld A, Dubcovsky J. Characterization of the maintained vegetative phase deletions from diploid wheat and their effect on *VRN2* and *FT* transcript levels. Mol Genet Genomics. 2010;283: 223–232.doi:10.1007/s00438-009-0510-2

49. Ream TS, Woods DP, Schwartz CJ, Sanabria CP, Mahoy JA, Walters EM, et al. Interaction of Photoperiod and Vernalization Determines Flowering Time of *Brachypodium distachyon*. Plant Physiol. 2014;164: 694–709. doi:10.1104/pp.113.232678

50. Woods DP, McKeown MA, Dong Y, Preston JC, Amasino RM. Evolution of *VRN2/Ghd7-Like* Genes in Vernalization-Mediated Repression of Grass Flowering. Plant Physiol. 2016;170: 2124–2135. doi:10.1104/pp.15.01279

51. Corbesier L, Vincent C, Jang S, Fornara F, Fan Q, Searle I, et al. FT Protein Movement Contributes to Long-Distance Signaling in Floral Induction of Arabidopsis. Science. 2007;316: 1030–1033. doi:10.1126/science.1141752

52. Tamaki S, Matsuo S, Wong HL, Yokoi S, Shimamoto K. Hd3a protein is a mobile flowering signal in rice. Science. 2007;316(5827):1033–6. doi:10.1126/science.1141753.

53. Cheng M-C, Kathare PK, Paik I, Huq E. Phytochrome Signaling Networks. Annu Rev Plant Biol. 2021;72: 217–244. doi:10.1146/annurev-arplant-080620-024221

54. Quail PH. Phytochrome Photosensory Signalling Networks. Nat Rev Mol Cell Biol. 2002;3: 85–93. doi:10.1038/nrm728

55. Möglich A, Yang X, Ayers RA, Moffat K. Structure and Function of Plant Photoreceptors. Annu Rev Plant Biol. 2010;61: 21–47. doi:10.1146/annurev-arplant-042809-112259

56. Mathews S. Evolutionary Studies Illuminate the Structural-Functional Model of Plant Phytochromes. Plant Cell. 2010;22: 4–16. doi:10.1105/tpc.109.072280

57. Chen A, Li C, Hu W, Lau MY, Lin H, Rockwell NC, et al. *PHYTOCHROME C* plays a major role in the acceleration of wheat flowering under long-day photoperiod. Proc Natl Acad Sci USA. 2014. doi:10.1073/pnas.1409795111

58. Woods DP, Ream TS, Minevich G, Hobert O, Amasino RM. PHYTOCHROME C is an essential light receptor for photoperiodic flowering in the temperate grass, *Brachypodium distachyon*. Genetics. 2014;198: 397–408. doi: 10.1534/genetics.114.166785

59. Pearce S, Kippes N, Chen A, Debernardi JM, Dubcovsky J. RNA-seq studies using wheat *PHYTOCHROME B* and *PHYTOCHROME C* mutants reveal shared and specific functions in the regulation of flowering and shade-avoidance pathways. BMC Plant Biol. BMC Plant Biology; 2016; 1–19. doi:10.1186/s12870-016-0831-3

60. Monte E, Alonso JM, Ecker JR, Zhang Y, Li X, Young J, et al. Isolation and characterization of *phyC* mutants in Arabidopsis reveals complex crosstalk between phytochrome signaling pathways. Plant Cell. 2003;15: 1962–1980.

61. Takano M, Inagaki N, Xie X, Yuzurihara N, Hihara F, Ishizuka T, et al. Distinct and cooperative functions of phytochromes A, B, and C in the control of deetiolation and flowering in rice. Plant Cell. 2005; 17: 3311–3325. doi:10.1105/tpc.105.035899

62. Kippes N, VanGessel C, Hamilton J, Akpinar A, Budak H, Dubcovsky J, et al. Effect of *phyB* and *phyC* loss-of-function mutations on the wheat transcriptome under short and long day photoperiods. BMC Plant Biology; 2020: 1–17. doi:10.1186/s12870-020-02506-0

63. Liu XL, Covington MF, Fankhauser C, Chory J, Wagner DR. ELF3 encodes a circadian clock–regulated nuclear protein that functions in an Arabidopsis PHYB signal transduction pathway. Plant Cell. 2001;13: 1293–1304.

64. Alvarez MA, Tranquilli G, Lewis S, Kippes N, Dubcovsky J. Genetic and physical mapping of the earliness. Funct Integr Genomics. 2016; 1–18. doi:10.1007/s10142-016-0490-3

65. Saito H, Ogiso-Tanaka E, Okumoto Y, Yoshitake Y, Izumi H, Yokoo T, et al. Ef7 encodes an ELF3-like protein and promotes rice flowering by negatively regulating the floral repressor gene *Ghd7* under both short-and long-day conditions. Plant and Cell Physiology. 2012;53: 717–728.doi:10.1093/pcp/pcs029

66. Dalmais M, Antelme S, Ho-Yue-Kuang S, Wang Y, Darracq O, d’Yvoire MB, et al. A TILLING Platform for Functional Genomics in *Brachypodium distachyon*. PLoS ONE. 2013;8: e65503–10. doi:10.1371/journal.pone.0065503

67. Woods D, Dong Y, Bouché F, Bednarek R, Rowe M, Ream T, et al. A florigen paralog is required for short-day vernalization in a pooid grass. Elife. 2019;8: 27. doi:10.7554/eLife.42153

68. Alvarez A, Li C, Lin H, Joe A, Padilla M, Woods DP and Dubcovsky J. *EARLY FLOWERING 3* interactions with *PHYTOCHROME B* and *PHOTOPERIOD1* are critical for the photoperiodic regulation of wheat heading time. PLoS Genetics. 2022

69. Nieto C, López-Salmerón V, Davière J-M, Prat S. ELF3-PIF4 Interaction Regulates Plant Growth Independently of the Evening Complex. Current Biology. 2015;25: 187–193. doi:10.1016/j.cub.2014.10.070

70. Liu XL, Covington MF, Fankhauser C, Chory J, Wanger DR. ELF3 encodes a circadian clock-regulated nuclear protein that functions in an Arabidopsis PHYB signal transduction pathway. Plant Cell. 2001;13: 1293–1304. doi:10.1105/tpc.13.6.1293

71. Yeom M, Kim H, Lim J, Shin A-Y, Hong S, Kim J-I, et al. How Do Phytochromes Transmit the Light Quality Information to the Circadian Clock in Arabidopsis? Molecular Plant. 2014;7: 1701–1704. doi:10.1093/mp/ssu086

72. Nieto C, López-Salmerón V, Davière J-M, Prat S. ELF3-PIF4 Interaction Regulates Plant Growth Independently of the Evening Complex. Current Biology. 2015;25: 187–193. doi:10.1016/j.cub.2014.10.070

73. Murakami M, Tago Y, Yamashino T, Mizuno T. Comparative Overviews of Clock-Associated Genes of *Arabidopsis thaliana* and *Oryza sativa*. Plant and Cell Physiology. 2006;48: 110–121. doi:10.1093/pcp/pcl043

74. Zhao J, Huang X, Ouyang X, Chen W, Du A, Zhu L, et al. *OsELF3-1*, an Ortholog of Arabidopsis *EARLY FLOWERING 3*, Regulates Rice Circadian Rhythm and Photoperiodic Flowering. PLoS ONE. 2012;7: e43705. doi:10.1371/journal.pone.0043705.s006

75. Itoh H, Tanaka Y, Izawa T. Genetic Relationship Between Phytochromes and *OsELF3-1* Reveals the Mode of Regulation for the Suppression of Phytochrome Signaling in Rice. Plant and Cell Physiology. 2019;60: 549–561. doi:10.1093/pcp/pcy225

76. R. Ishikawa et al. *Phytochrome B* regulates *Heading date 1* (*Hd1*)-mediated expression of rice florigen *Hd3a* and critical day length in rice. Mol. Genet. Genomics. 2011; 285:461–470. doi: 10.1007/s00438-011-0621-4

77. Koo B-H, Yoo S-C, Park J-W, Kwon C-T, Lee B-D, An G, et al. Natural variation in *OsPRR37* regulates heading date and contributes to rice cultivation at a wide range of latitudes. Molecular Plant. 2013;6: 1877–1888. doi:10.1093/mp/sst088

78. Murphy RL, Klein RR, Morishige DT, Brady JA, Rooney WL, Miller FR, et al. Coincident light and clock regulation of pseudoresponse regulator protein 37 (*PRR37*) controls photoperiodic flowering in sorghum. Proc Natl Acad Sci USA. 2011;108: 16469–16474. doi:10.1073/pnas.1106212108

79. Gordon SP, Contreras-Moreira B, Woods DP, Marais Des DL, Burgess D, Shu S, et al. Extensive gene content variation in the *Brachypodium distachyon* pan-genome correlates with population structure. Nature Communications. 2017;8: 2184. doi: 10.1038/s41467-017-02292-8.

80. Woods DP, Ream TS, Bouché F, Lee J, Thrower N, Wilkerson C, et al. Establishment of a vernalization requirement in *Brachypodium distachyon* requires *REPRESSOR OF VERNALIZATION1*. Proc Natl Acad Sci USA. 2017;114: 6623–6628. doi:10.1073/pnas.1700536114

81. Dubcovsky J, Loukoianov A, Fu D, Valarik M, Sanchez A, Yan L. Effect of Photoperiod on the Regulation of Wheat Vernalization Genes *VRN1* and *VRN2*. Plant Mol Biol. 2006;60: 469–480. doi:10.1007/s11103-005-4814-2

82. Gawronski P, Ariyadasa R, Himmelbach A, Poursarebani N, Kilian B, Stein N, et al. A distorted circadian clock causes early flowering and temperature-dependent variation in spike development in the *Eps-3A^m^* mutant of einkorn wheat. Genetics. 2014;196(4):1253–1266. doi: 10.1534/genetics.113.158444

83. De Mendiburu R-project org package agricolae accessed 25 July F, 2012. Agricolae: Statistical procedures for agricultural research. R package version 1.1-3. Comprehensive R Arch.

84. Shaw LM, Lyu B, Turner R, Li C, Chen F, Han X, Fu D, Dubcovsky J. *FLOWERING LOCUS T2* regulates spike development and fertility in temperate cereals. J Exp Bot. 2019; 70:193–204. doi:10.1093/jxb/ery35

