## Supplemental Table 1 for "PHYTOCHROME C regulation of *PHOTOPERIOD1* is mediated by *EARLY FLOWERING 3* in *Brachypodium distachyon*"

| Purpose | Primers | Sequence | NEB enzyme | Reference |
|---|---|---|---|---|
| qPCR | qUBC18-F | GTCACCCGCAATGACTGTAAGTTC | | Ream et al., 2014 |
| | qUBC18-R | TTGTCTTGCGGACGTTGCTTTG | | |
| | qFT1-F | TTCGGGAACAGGAACGTGTCCAAC | | |
| | qFT1-R | AGCATCTGGGTCTACCATCACGAG | | |
| | qVRN1-F | GCTCTGCAGAAGGAACTTGTGG | | |
| | qVRN1-R | CTAGTTTGCGGGTGTGTTTGCTC | | |
| | qVRN2-F | ATGCATGAGAGAGAGGCGAAGG | | |
| | qVRN2-R | TCGTAGCGGATCTGCTTCTCGTAG | | |
| | qPPD1-F2 | CTATGCCGTCGCTTGAGTTG | | This paper |
| | qPPD1-R2 | TGCCGCCTTGATTGGAAACC | | |
| | qCO1-F | AGAGTGGTTATGGGCTTGGA | | |
| | qCO1-R | CTATCACCGTATTGTCTGGG | | |
| | qCO2-F | GGCAAGTGAGGAACAGGAAAG | | |
| | qCO2-R | TAGGCTCCACTGGTTGTTAGG | | |
| Fine Mapping and genotyping | dCaps16020-F | CCGTCTCCAGATTATACCTATCCG | Hpy188I | This paper |
| | dCaps16020-R | GAGTGTGATTTCAGCCCTTG | | |
| | pdd1_F | CAACCGGAGATGGTGGAAATG | Hpy166II | |
| | pdd1_R | GGACACTGATATTATGTTGTGC | | |
| | dCaps17600-F | AAAGTCCTGCTGGCGCTCGTG | BsrDI | |
| | dCaps17600-R | TGACGCCGTGCTCCGGCTCTGCAA | | |
| | elf3-F | CACCCATGCCTCCAATGTACTTCCC | Hpy166II | Bouché et al. 2022 |
| | elf3-R | GGTGGTTTCAGCTTCTGCAGGTGAA | | |

Table S1. Primers used in this study
